# Immune-competent apical-out airway organoids reveal distinct antiviral strategies of macrophages, neutrophils, and monocytes during influenza infection

**DOI:** 10.64898/2026.06.03.729766

**Authors:** Martin Beukema, Jacqueline J. de Vries-Idema, Nilima Dinesh Kumar, Shuran Gong, Nils Doesburg, Anke L.W. Huckriede, Barbro N. Melgert

## Abstract

The respiratory mucosa is the primary entry site for influenza virus and early antiviral immunity is governed there by interactions between epithelial cells and innate immune cells. However, mechanistic insight into these interactions in a human context is limited. Here, we established an immune-competent apical-out human airway organoid model that enables direct epithelial infection and controlled integration of innate immune cells. The organoids recapitulate key features of the human upper airway epithelium and support productive influenza virus replication. Using defined co-cultures, we uncovered distinct innate immune functions: macrophages suppress viral replication and restrict epithelial spread via production of type I and III interferons; neutrophils reduce extracellular virus levels without limiting epithelial infection, consistent with antiviral clearance mechanisms independent of interferon signaling; and monocytes exert modest, transient antiviral effects. Together, we use a human-relevant platform to define how distinct innate immune cells shape mucosal immunity at the respiratory epithelium, with direct implications for antiviral and vaccine development.

**Graphical Abstract:** 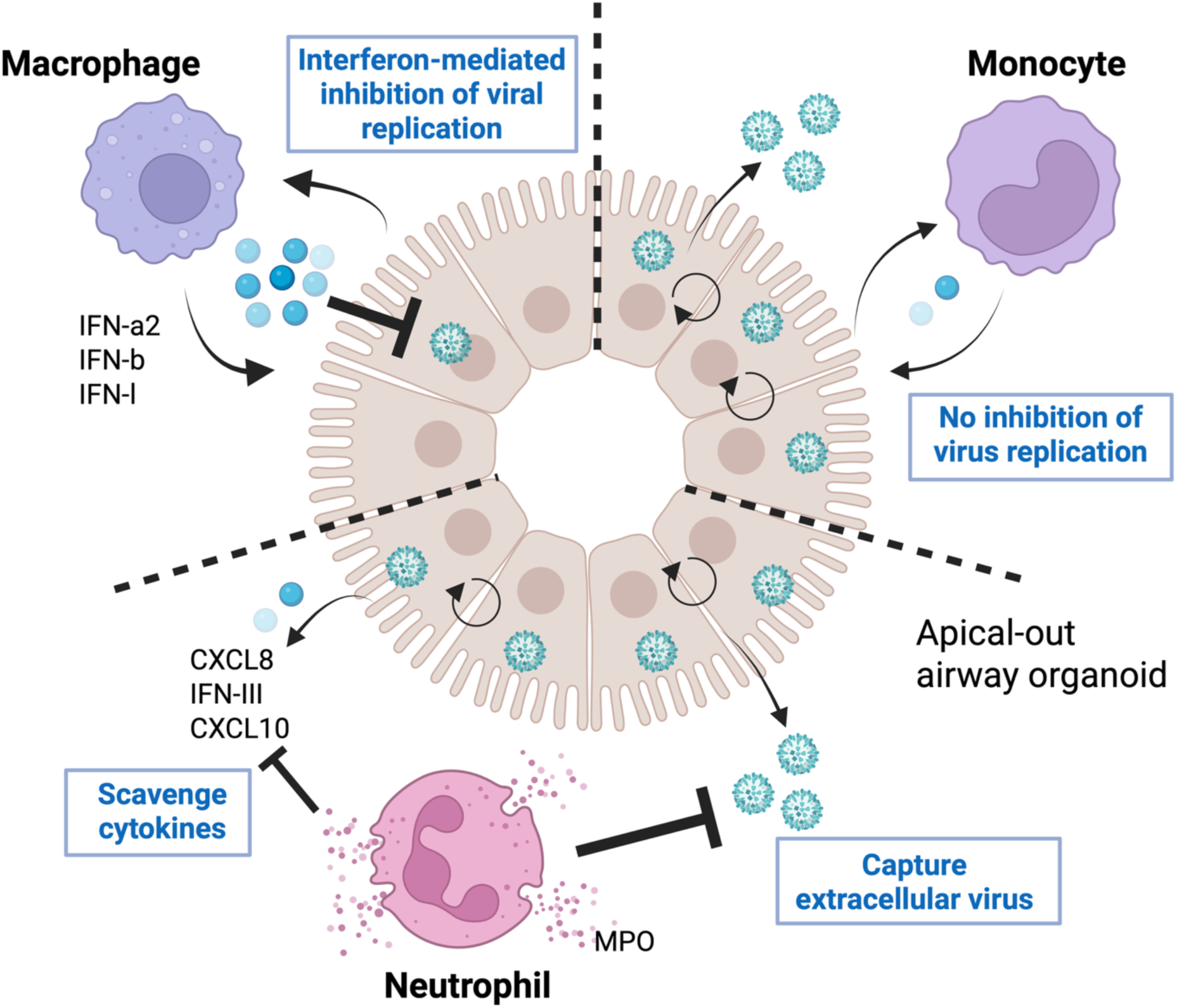

## Introduction

Influenza virus continues to pose a significant global health burden, driving recurrent outbreaks of respiratory illness that range from mild upper airway symptoms to life-threatening complications (1). Although advances in vaccination strategies have been made over the years, influenza continues to be a major health concern (2). This limitation stems in part from the incomplete understanding of early human immune responses to influenza infection at the mucosa, the primary site of infection (3,4). This limits the development of effective protective strategies against infection with influenza virus.

The mucosa of the upper respiratory tract serves as the primary entry point for human influenza viruses. Infection begins with viral attachment to α2-6-sialic acids and replication in epithelial cells lining the upper airways (5). This triggers a rapid and coordinated antiviral response, characterized by the release of cytokines and chemokines that orchestrate immune cell recruitment (6). Studies in animals have shown that infection outcomes depend on dynamic interactions between epithelial cells and (recruited) innate immune cells, including macrophages, neutrophils, and monocytes (7–9). Upon infection of epithelium, tissue-resident macrophages act rapidly as immunological sentinels, detecting and phagocytosing viral components, initiating inflammatory signaling, and coordinating recruitment of additional immune cells (10). Neutrophils are among the first recruited responders, infiltrating infected tissues from the blood circulation and deploying antimicrobial strategies such as phagocytosis, degranulation, and the formation of neutrophil extracellular traps (11). After neutrophils, monocytes migrate into inflamed mucosal sites, differentiating into macrophages and/or dendritic cells that contribute to cytokine production and antigen presentation and coordination of tissue repair (10). Yet, understanding how these innate immune cells interact with the respiratory epithelium during influenza virus infection and contribute to antiviral immunity in a human-relevant context remains a critical challenge (12). Addressing this gap may enable the identification of cellular targets for protective strategies.

Although animal models have provided valuable insights into host–influenza virus interactions, their inherent complexity and species-specific differences limit the resolution and translational relevance of studying epithelial–immune dynamics during human influenza virus infection (12,13). Airway organoids have emerged as more physiologically relevant tools for modeling the cellular architecture and complexity of human respiratory epithelium, offering a promising platform for investigating host–virus interactions (14–16). However, most studies have relied on extracellular matrix (ECM)-embedded organoids, which form a polarized apicobasal epithelium surrounding a central lumen, commonly referred to as apical-in organoids. These models pose technical challenges for viral infection studies as the apical surface is not readily available for exogenously added virus. Accordingly, infection requires microinjection of the virus into the lumen (17) or mechanical disruption of organoids with subsequent virus addition (18). Recent advances in organoid technology have enabled the development of apical-out (AO) airway organoids, in which the epithelial surface is oriented outward, allowing direct access for viruses and providing a more physiologically representative model for studying viral entry and replication (19). However, to date, AO airway organoids have primarily been used as tool for virus infection. They have not been characterized for epithelial immune responses or used for incorporation of innate immune cells to investigate early antiviral responses (12).

In this study, we aimed to compare how macrophages, neutrophils, and monocytes shape early innate immune responses to influenza A virus (IAV) infection in human airway epithelium within a physiologically relevant context. To test this, We established and characterized a human-relevant, immune-competent airway organoid (AO) system that recapitulates key features of the upper airway epithelium. Using this platform, we systematically investigate how distinct innate immune cell populations influence IAV infection outcomes in human airway epithelial cells.

## Material and methods

### Isolation of airway basal cells

Residual tracheal and main stem bronchial tissues were obtained post-mortem from lung transplant donors following organ retrieval. Donor selection adhered to Eurotransplant guidelines, requiring the absence of primary lung disease (e.g., asthma, COPD) and a smoking history of no more than 20 pack-years. Until epithelial cell isolation, tissues were stored at 10°C in Krebs buffer (117.5 mM NaCl, 5.6 mM KCl, 1.18 mM MgSO₄·7H₂O, 1.28 mM NaH₂PO₄·H₂O, 2.52 mM CaCl₂·2H₂O, 25 mM NaHCO₃, 5.55 mM D-glucose·H₂O in sterile H₂O).

Following excision, tissues were trimmed, rinsed, and incubated for 2 hours at 37 °C in calcium-and magnesium-free Hanks’ Balanced Salt Solution (HBSS; Lonza, #14175) supplemented with 100 IU/ml penicillin/streptomycin (Gibco, #15140-122) and 0.18 mg/mL protease XIV (Sigma, #P5147, 4P8). Epithelial cells were gently scraped from the luminal surface of the trachea, washed twice, and seeded into six-well plates (Greiner, #657970) pre-coated for 2–6 hours with a mixture of 30 μg/mL PureCol (Advanced BioMatrix, #5005-B), 10 μg/mL fibronectin (Sigma, #F0895), and 10 μg/mL bovine serum albumin (BSA; Sigma, #A2153) in Phosphate Buffered Saline (PBS, Gibco, #1404-091). Cells were cultured in keratinocyte serum-free medium (KSFM; Gibco, #17005-034) supplemented with 0.2 ng/mL epidermal growth factor (EGF; Gibco, #17005-075), 25 μg/mL bovine pituitary extract (BPE; Gibco, #17005-075), 1 μM isoproterenol (Sigma, #I-6504), 200 U/mL penicillin, and 200 μg/mL streptomycin. Ciprofloxacin (10μg/ml, Sigma, #17850) was added during the first week of culture to prevent mycoplasma contamination. Upon reaching ∼70% confluency, cells were dissociated using an animal component-free cell dissociation kit (Stemcell, #05426) in combination with 0.2 g/L EDTA (Lonza, #BE17-161E), and expanded in Ex-plus medium (Stemcell, #05040). Once ∼90% confluency was reached, airway cells were trypsinized (Stemcell, #05426) and cryopreserved in Bambanker serum-free freezing medium (Nippon Genetics, #BBD01).

### Generation of apical-out airway organoids

Airway organoids with AO polarization were generated with PneumaCult^TM^ AO airway organoid medium (Stemcell, #100-0620) according to the manufacturer’s instructions. In short, cryo-preserved basal cells were thawed and expanded for 3 days in Ex-plus medium (Stemcell, #05040) to reach ∼60% confluency. Next, cells were trypsinized using the animal component-free cell dissociation kit (ACK; Stemcell, # 05426,) and seeded (120.000 cells) in AggreWell™400 24-well plates (Stemcell, #34450), pretreated with anti-adherence solution (Stemcell, #07010). Cells were cultured for two days in the Aggrewells with AO airway organoid medium to generate aggregates. After two days, the aggregates were transferred to a 24-wells plate and differentiated to AO airway organoids for another 12 days. AO airway organoid medium was replaced twice per week.

### PBMC, neutrophil, monocyte, and macrophage isolation

Human neutrophils and peripheral blood mononuclear cells (PBMCs) were isolated from buffy coats of healthy donors, obtained from the Dutch Blood Bank (Sanquin, Nijmegen, NL), according to a previous study (20). Polymorphonuclear cells (PMNs) and mononuclear cells were separated by density gradient centrifugation (800 × g, 30 min) using Ficoll Histopaque (Cytiva, #17144003). PBMCs were collected as previously described (21).

Neutrophils were isolated from peripheral blood (n = 6) using density gradient centrifugation. To minimize pre-activation, all buffers and media were pre-cooled on ice throughout the isolation procedure. Following centrifugation over Ficoll, all layers above the granulocyte–erythrocyte interface were carefully removed. The granulocyte–erythrocyte fraction was transferred to a fresh tube, and residual erythrocytes were lysed using ice-cold ACK lysing buffer (Gibco Life Technologies, NY, USA; #A10492-01) at a 2.5:1 (v/v) buffer-to-cell suspension ratio. During lysis, tubes were gently inverted every 30 seconds. Lysis was terminated once the suspension darkened and viscosity decreased, after which Hank’s Balanced Salt Solution (HBSS; Ca²⁺/Mg²⁺-free; Gibco Life Technologies, Paisley, UK) supplemented with 5% fetal bovine serum (FBS; Life Science Production, Bedfordshire, UK) was added to a final volume of 50 mL. Cells were centrifuged at 200 × g for 3 minutes, and the supernatant was discarded. If erythrocyte contamination remained, the lysis step was repeated. Purified neutrophils were resuspended in ice-cold complete culture medium, and cell viability was assessed by Trypan Blue exclusion (Merck Life Science, Gillingham, UK) and confirmed by flow cytometry.

PBMC-derived monocytes (n = 6) were isolated using the EasySep™ Human Monocyte Negative Selection Kit (Stemcell Technologies, #19359). Monocyte-derived macrophages (n = 5) were generated by culturing monocytes in the presence of 100 ng/mL recombinant macrophage colony-stimulating factor (BioLegend, #574804) for 6 days (22). On day 6, macrophages were detached using 1× citrate buffer (1.35 M KCl, 174.38 mM Sodium Citrate in PBS).

### Influenza virus infection of airway organoid-immune cell co-cultures

Upon 14 days of differentiation, AO airway organoids were infected with the influenza virus. Prior to infection, epithelial cell numbers in AO airway organoids were quantified by dissociating organoids from a single well using ACK dissociation solution (Stemcell Technologies). Then organoids containing approximately 300.000 epithelial cells were incubated for 1 hour with the influenza A/Wisconsin/588/2019 (H1N1)pdm09 virus (Medicines and Healthcare products Regulatory Agency [MHRA] repository, UK) at a multiplicity of infection (MOI) of 0.1, 0.01 or 0.001 in AO airway organoid medium without heparin. After 1 hour, organoids were washed with advanced DMEM/F-12 medium (Gibco, #12634-010) supplemented with 1 M HEPES (Gibco, #15630-080), 500 μl 1x glutamax (Gibco, #35050061), 100 IU/mL penicillin and 100 μg/mL streptomycin penicillin/streptomycin (Gibco, #15140-122). In co-culture experiments, either 120.000, 60.000 or 30.000 immune cells were cultured with the infected organoids in AO airway organoid medium without heparin (#100-0620, Stemcell). Cells from one epithelial donor showing average levels of virus-induced cytokine levels (see Legendplex below) were used for co-culture experiments. For IFN inhibition assays, cultures were incubated with 400 nM ruxolitinib (JAK1/2 inhibitor) (22). For phagocytosis inhibition assays, macrophages or neutrophils were pretreated for 30 min with 5 μM Cytochalasin D (23). Next, the cells were washed twice with PBS before culturing with infected AO airway organoids.

At 3, 6, 12, 24, and 48 hours post infection (hpi), supernatants were collected for virus and cytokine/chemokine quantifications. The organoids were fixed in 500 µl 4% paraformaldehyde (PFA, Sigma Aldrich, #158127-500G), washed, and incubated in PBS (Gibco) at 4°C until immunofluorescent staining.

### Microscopic characterization of epithelial cell types and influenza infection

AO airway organoids were characterized for the presence of basal cells (p63), goblet cells (MUC5a), club cells (CC10), ciliated cells (acetylated α tubulin), alveolar type 2 (pro surfactant protein C), and influenza infection receptors α2-3-sialic acid and α2-6-sialic acid. Furthermore, organoids were stained for nucleoprotein to visualize influenza infection. The staining of PFA-fixed organoid was performed in 1.5 ml Eppendorf tubes. First, organoids were permeabilized with PBS (Gibco) solution containing 0.5% v/v Triton-X100 (Sigma Aldrich, T-8787) for 10 min at RT. Next, the organoids were blocked for 15 min. with 5% w/v bovine serum albumin (BSA) containing 5% v/v normal human serum in PBS. After blocking, organoids were incubated overnight with primary antibody (Table 1) in PBS containing 2% w/v BSA and 1% v/v normal human serum at 4°C. The next day, organoids were blocked with 5% w/v BSA and 5% v/v normal goat serum or normal rabbit serum in PBS, followed by a 2-hour incubation with a 1:300 dilution of the respective secondary antibody (Table 1) and a 1:400 dilution of phalloidin in PBS. Next, organoids were suspended in 50 µl Prolong Gold antifade reagent with DAPI (Invitrogen, #p36931), before being added to a glass microscope slide (Knittel-Glaeser). Between the separate incubation steps, organoids were washed twice with PBS. Images were acquired using a Ti2 microscope (Nikon, Japan). Scale bars were added using FIJI software.

**Table 1.**
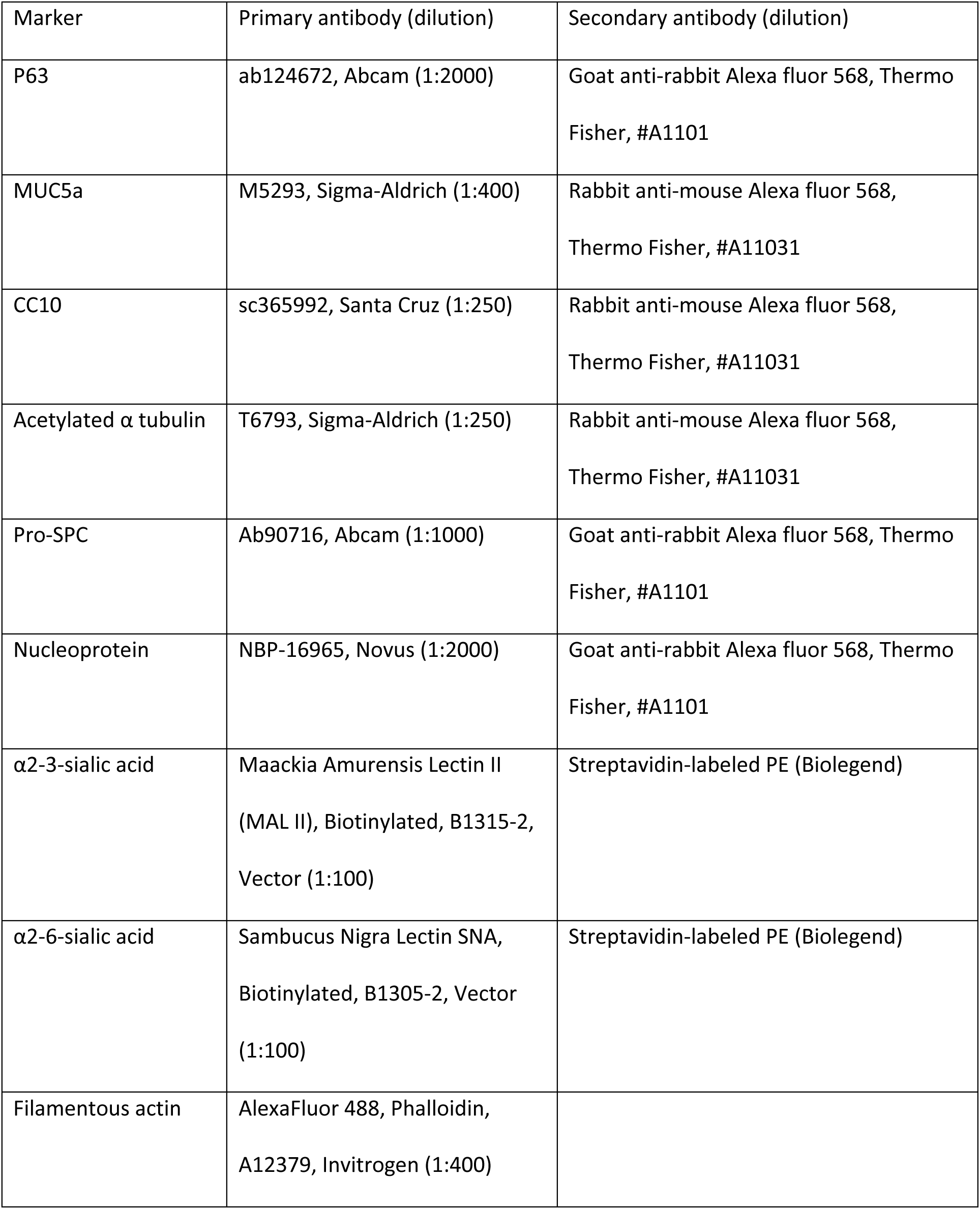
Primary and secondary antibodies for immunofluorescent staining.

| Marker | Primary antibody (dilution) | Secondary antibody (dilution) |
| --- | --- | --- |
| P63 | ab124672, Abcam (1:2000) | Goat anti-rabbit Alexa fluor 568, Thermo Fisher, #A1101 |
| MUC5a | M5293, Sigma-Aldrich (1:400) | Rabbit anti-mouse Alexa fluor 568, Thermo Fisher, #A11031 |
| CC10 | sc365992, Santa Cruz (1:250) | Rabbit anti-mouse Alexa fluor 568, Thermo Fisher, #A11031 |
| Acetylated $\alpha$ tubulin | T6793, Sigma-Aldrich (1:250) | Rabbit anti-mouse Alexa fluor 568, Thermo Fisher, #A11031 |
| Pro-SPC | Ab90716, Abcam (1:1000) | Goat anti-rabbit Alexa fluor 568, Thermo Fisher, #A1101 |
| Nucleoprotein | NBP-16965, Novus (1:2000) | Goat anti-rabbit Alexa fluor 568, Thermo Fisher, #A1101 |
| α2-3-sialic acid | Maackia Amurensis Lectin II (MAL II), Biotinylated, B1315-2, Vector (1:100) | Streptavidin-labeled PE (Biolegend) |
| α2-6-sialic acid | Sambucus Nigra Lectin SNA, Biotinylated, B1305-2, Vector (1:100) | Streptavidin-labeled PE (Biolegend) |
| Filamentous actin | AlexaFluor 488, Phalloidin, A12379, Invitrogen (1:400) |  |

### Virus titrations

Virus titers were quantified using a tissue culture infectious dose 50 (TCID50)-based hemagglutination (HA) assay as described before (24). Madin-Darby Canine Kidney (MDCK) cells were seeded into flat-bottom 96-well plates and cultured overnight in serum-free EPISERF medium (Gibco, #10732-022). The following day, confluent monolayers were inoculated with serial 2-fold dilutions of virus-containing supernatants prepared in EPISERF medium. A volume of 225 µL was added to each well. After 1 hour of incubation at 37°C and 5% CO₂, the inoculum was removed, and cells were washed once with PBS (Gibco). Subsequently, cells were incubated for 72 hours in EPISERF medium supplemented with 6 µg/mL TPCK-treated trypsin (Sigma Aldrich, # T1426). At the end of the incubation period, 100 µL of the supernatant from each well was transferred to a V-bottom 96-well plate. To each well, 50 µL of a 1.5% suspension of guinea pig erythrocytes was added. After 2 hours of incubation at room temperature, wells were scored for hemagglutination activity. The TCID50 was calculated using the Spearman–Kärber method (25).

### Multiplex cytokine analysis

The cytokines and chemokines IL-1β, IL-6, CXCL8 (IL-8), IL-10, IL-12p70, IFN-α2, IFN-β, IFN-λ1 (IL-29), IFN-λ2 (IL-28a), IFN-γ, TNF-α, CXCL10 (IP-10), and GM-CSF were determined using the LEGENDplex^TM^ Human Anti-Virus Response Panel 1 (13-plex) w/VbP V02 kit (Biolegend, # 741270).

### Myeloperoxidase ELISA

To determine the level of myeloperoxidase (MPO) in the supernatants of cultures, the LEGEND MAX™ Human Myeloperoxidase ELISA Kit (Biolegend, # 440007) was employed according to the manufacturer’s instructions. Culture supernatant samples were diluted 1:10 with Assay Buffer B in this assay.

### Statistics

Statistical analyses were performed using GraphPad Prism version 10 for Windows, GraphPad Software, Boston, Massachusetts USA, www.graphpad.com. Statistical differences in virus titers were measured by comparing geometric mean in log-transformed parametric analysis. Nonparametric tests were used for the other parameters. The type of analysis used is indicated in the respective figure legend.

## Results

### Apical-out airway organoids represent a physiologically relevant model of the upper airways for viral entry and replication

To model the early stages of influenza infection in a physiologically relevant context, we employed AO airway organoids (**Fig. 1**). This model features AO polarization as evidenced by the presence of differentiated epithelial cell types at the outer surface of the organoids, making them accessible to viral infection (**Fig. 1**). The AO airway organoid model predominantly consists of ciliated cells (**Fig. 1A**) and some goblet cells (**Fig. 1B**), but no club cells (**Suppl. Fig. 1A**), or alveolar type 2 cells (**Suppl. Fig. 1B**). Basal cells were also absent, suggesting full differentiation of basal to ciliated or goblet cells (**Fig. 1C**). Furthermore, the AO airway organoids mainly expressed α2-6-sialic acid receptors (**Fig. 1E**) and only few α2-3-sialic acid receptors (**Fig. 1D**). The observed epithelial composition and sialic acid expression profiles correspond with the human situation, where the upper respiratory tract is largely composed of ciliated cells that highly express α2-6-sialic acid receptors (5, 26).

**Figure 1.**
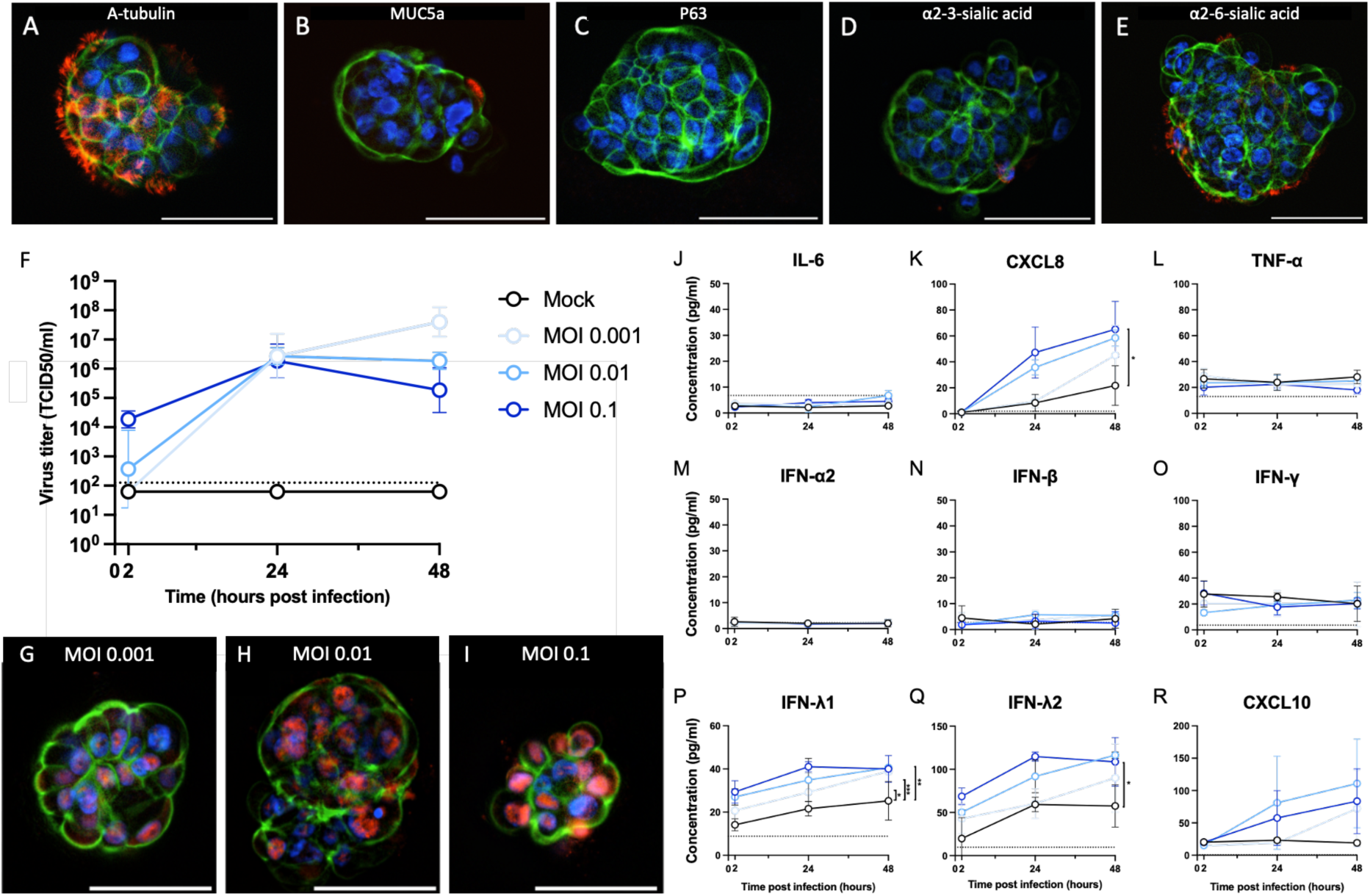
Virus-host interactions between apical out organoids and the influenza virus. The organoids were fluorescently stained to detect the presence of ciliated epithelial cells (A; α-tubulin), goblet cells (B; MUC5a), basal cells (C; p63), α2-3-sialic acid (D; Maackia Amurensis Lectin II), and α2-6-sialic acid (E; Sambucus Nigra Lectin). Blue dye indicates nuclear staining (DAPI). Green dye indicates filamentous actin (Phalloidin). Influenza virus titers were determined at 2, 24, and 48 hours post-infection at the indicated MOIs (F; N = 3). At 24 hours post-infection, organoids infected with an MOI of 0.001 (G), MOI of 0.01 (H), and MOI of 0.1 (I) were stained for influenza virus (red dye, nucleoprotein; N=3). Blue dye indicates nuclear staining (DAPI). Green dye indicates filamentous actin (Phalloidin). Cytokines IL-6 (J), CXCL8 (K), TNF-α (L), IFN-α2 (M), IFN-β (N), IFN-γ (O), IFN-λ1 (P), IFN-λ2/3 (Q), and CXCL10 (R) were determined in supernatants of infected organoids at 2, 24, and 48 hours post-infection. (N = 3). Virus titers and cytokine data are represented as geometric mean ± SD and mean ± SD, respectively. Statistical differences in the area under the curve (AUC) of viral titers and cytokine responses between MOI conditions were assessed using the log-normalized repeated measures one-way ANOVA followed Tukey posthoc test and Friedman test followed by Dunn’s post hoc test, respectively (* = *p* < 0.05, ** = *p* < 0.01, *** = *p* < 0.001). Fluorescent microscopy pictures were taken at 400x magnification. Scalebar = 50 μM.

Next, we assessed infectivity and replication dynamics of influenza A/Wisconsin/588/2019 (H1N1)pdm09 in this airway organoid model. Organoids were infected with three different MOIs: 0.001, 0.01, and 0.1 (**Fig. 1F**). Surprisingly, organoids infected with the lowest viral dose (MOI 0.001) produced the highest viral titers at both 24 and 48 hours post-infection, followed by those infected at MOI 0.01 and MOI 0.1, respectively. Despite lower overall viral output, organoids infected at higher MOIs (0.01 and 0.1) displayed more intense staining for influenza nucleoprotein (NP) (**Fig. 1H** and **1I**) compared to those infected at MOI 0.001 (**Fig. 1G**), indicating greater initial viral uptake and protein expression. Notably, infection at an MOI of 0.1 led to structural disintegration of the organoids at 24 hours post-infection.

To investigate epithelial responses to influenza A infection, chemokine/cytokine secretion induced by infection of AO airway organoids was measured in culture supernatants (**Fig. 1J-R**). In the absence of infection (Mock), organoids exhibited only modest CXCL8 (**Fig. 3K**), IFNλ-1 (**Fig. 3P**), and IFN-λ2 (**Fig. 3Q**) production over time. Upon influenza A infection, however, the organoids produced significantly higher levels of these chemokines and cytokines. Infection also induced a trend towards increased CXCL10 production (**Fig. 1R**). These cytokines were secreted in a MOI-dependent manner, with higher viral doses resulting in more cytokine secretion. Although an MOI of 0.001 resulted in high viral replication (**Fig. 1F**), cytokine levels remained close to those in uninfected organoids. In contrast, infection at an MOI of 0.01 sustained viral replication up to 48 hours post-infection and elicited a robust cytokine response. Therefore, the MOI of 0.01 was selected for subsequent co-culture experiments. Overall, these results confirm that apical out-oriented airway epithelium in organoids supports productive influenza infection and can serve as a physiologically relevant model for viral entry and replication. This model therefore provided a robust platform for subsequent co-culture experiments with immune cells to study host-pathogen interactions in a controlled, mucosal tissue-like environment.

### Macrophages exert potent antiviral effects against influenza infection in AO airway organoids

Following characterization of viral replication in AO airway organoids cultured alone, we next investigated effects of macrophages (**Fig. 2**), neutrophils (**Fig. 4**), or monocytes (**Fig. 6**) on organoids during influenza A virus infection. To resemble *in vivo* infection of the epithelium, organoids were pre-incubated for 1 hour with influenza virus first before immune cells were added. Unbound virus was removed before addition of immune cells.

**Figure 2.**
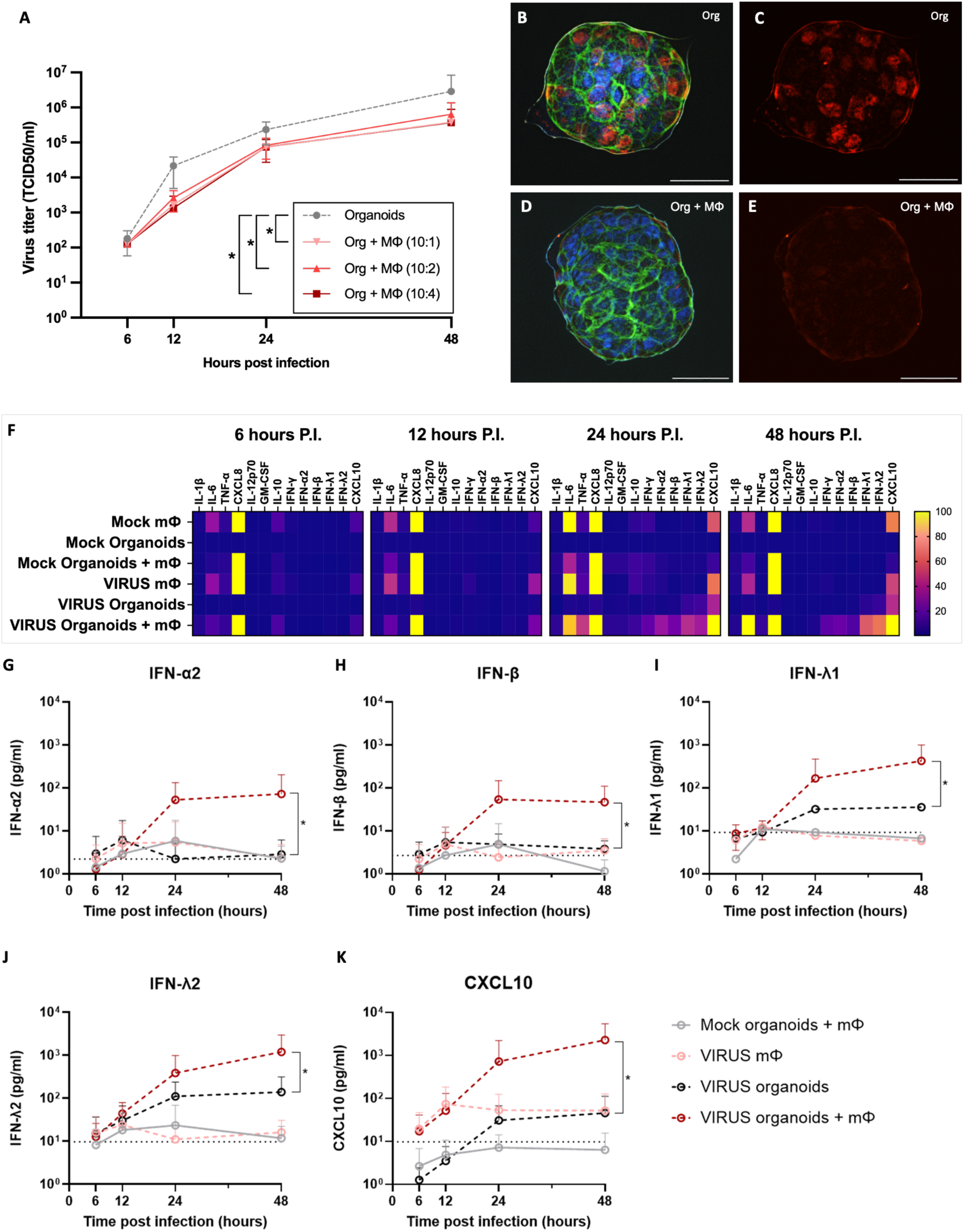
Impact of macrophages on influenza virus infection in AO airway organoids. Influenza virus titers were quantified from co-cultures of infected organoids (MOI 0.01) with macrophages (A). At 24 hours post-infection, organoids were stained to visualize influenza virus dissemination. Infected organoids cultured without (B, C) or with macrophages (D, E) were stained for influenza virus nucleoprotein (red), nuclei (DAPI, blue), and filamentous actin (Phalloidin, green). Cytokine levels were measured from culture supernatants at 6, 12, 24, and 48 hours post-infection and plotted as fold-change relative to mock-infected organoids. Specifically, levels of antiviral cytokines IFN-α2 (G), IFN-β (H), IFN-λ1 (I), IFN-λ2 (J), and the IFN-inducible chemokine CXCL10 (K) are shown. Fluorescent microscopy pictures were taken at 400x magnification. Virus titers and cytokine data are represented as geometric mean ± SD and mean ± SD, respectively. Statistical differences in the area under the curve (AUC) of viral titers and cytokine responses between conditions were assessed using the log-normalized repeated measures one-way ANOVA followed Tukey posthoc test and Friedman test followed by Dunn’s post hoc test, respectively (* = *p* < 0.05). Scalebar = 50 μM. mФ = macrophages; P.I. = post infection.

Co-culturing organoids with macrophages significantly suppressed viral replication (>10-fold) in AO airway organoids over a 48-hour period, regardless of the macrophage number used (**Fig. 2A**). In the absence of macrophages, influenza virus could spread extensively throughout the organoids (**Fig. 2B,C**). In contrast, macrophage presence markedly restricted viral dissemination throughout the organoids (**Fig. 2D,E**), with influenza virus confined to the organoid periphery. Viral replication only occurred in epithelial cells, and not in virus-exposed macrophages (data not shown). These results indicate that macrophages limit viral replication in epithelial cells and prevent the spread of the virus to neighboring epithelial cells.

To identify which factors modulate macrophage-mediated viral control in AO airway organoids, we quantified key antiviral cytokines secreted during infection (**Fig. 2F**). Both macrophage monocultures and co-cultures containing macrophages showed a rapid and sustained increase in CXCL8 and IL-6 secretion, persisting up to 48 hours after infection. In response to infection, macrophages markedly elevated cytokines associated with proinflammatory macrophages (IL-6, TNF-α, and IFN-γ) both in absence and presence of organoids. Co-culturing macrophages with infected organoids further amplified these responses, whereas the anti-inflammatory cytokine IL-10 remained at moderate levels. Moreover, Type I and III interferons (IFNs) were also significantly higher in all macrophage-containing cultures, with the highest levels observed in co-cultures of macrophages and infected organoids. These included IFN-α2 (**Fig. 2I**), IFN-β (**Fig. 2H**), IFN-λ1 (**Fig. 2I**), IFN-λ2 (**Fig. 2J**), and CXCL10 (**Fig. 2K**), all exceeding levels in infected organoids alone. Elevated cytokine secretion persisted for up to 48 hours, though responses were attenuated when macrophage numbers were lower (**Suppl. Fig. 2A**). Collectively, these findings suggest that macrophages and epithelial cells synergistically enhance cytokine secretion during influenza virus infection, particularly of type I and type III IFNs.

To determine whether the macrophage-mediated reduction in influenza virus replication was driven by enhanced interferon (IFN) signaling, phagocytosis, or both, influenza-infected organoid–macrophage co-cultures were treated with the JAK1/2 inhibitor ruxolitinib, the phagocytosis inhibitor cytochalasin D, or a combination of both (**Fig. 3**). Treatment with ruxolitinib markedly reduced IFN secretion in both organoid mono-cultures and organoid–macrophage co-cultures during infection (**Fig. 3B)**. This reduction in IFN signaling was accompanied by an increase in viral titers, confirming that IFN responses are critical for limiting virus replication in macrophage-organoid cultures. Pretreatment of macrophages with cytochalasin D further reduced viral titers in culture supernatants (**Fig. 3A**). This effect coincided with strongly elevated levels of type I and type III IFNs (**Fig. 3B**). Combined treatment with cytochalasin D and ruxolitinib dampened the cytochalasin D–induced increase in type I IFNs (**Fig. 3B**) and showed a concomitant trend toward higher viral titers. Collectively, these data indicate that macrophages restrict influenza virus replication in airway organoids primarily through the induction of type I and type III IFNs, rather than through direct phagocytic clearance of viral particles.

**Figure 3.**
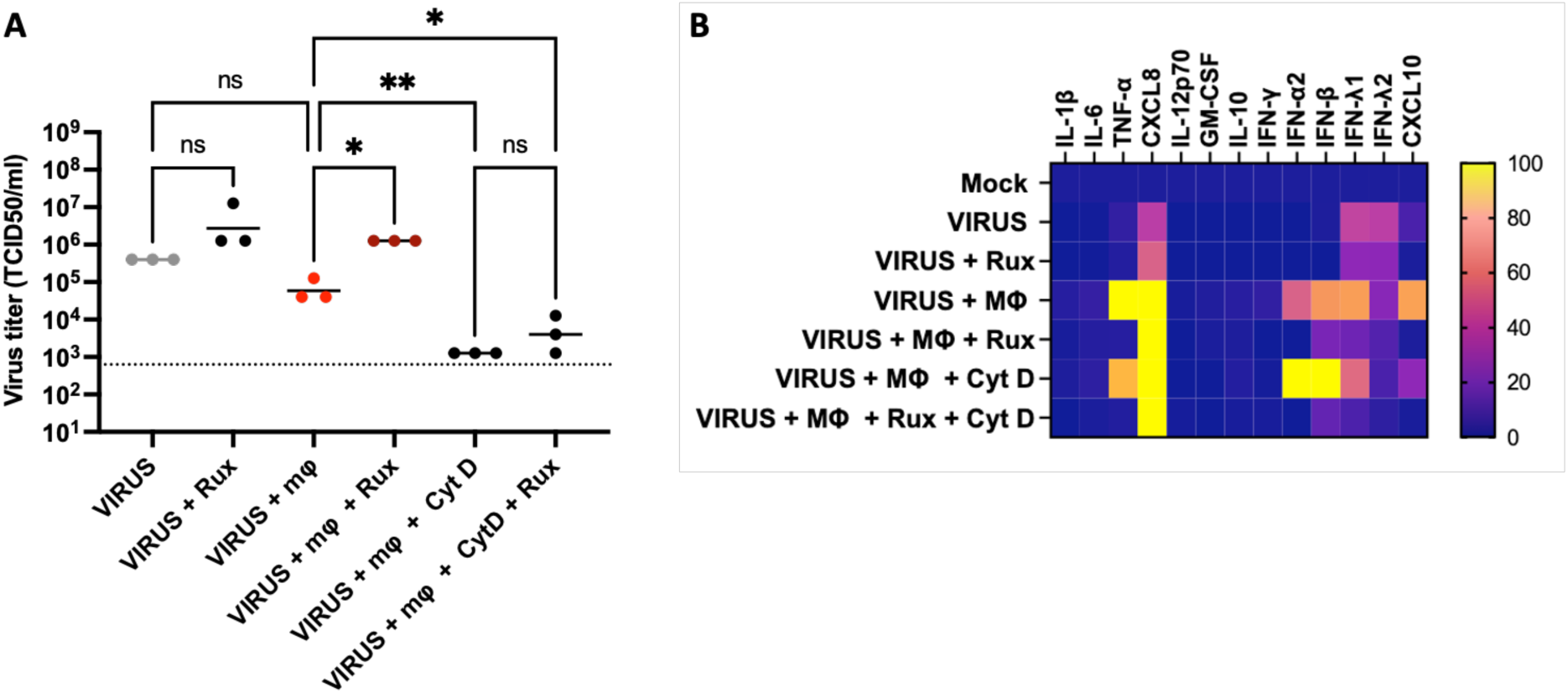
Factors underlying the antiviral role of macrophages. Influenza virus titers were quantified from infected AO airway organoid cultures (VIRUS), and co-cultures between macrophages and infected AO airway organoid (A). To inhibit IFN signaling, the JAK1/2 inhibitor ruxolitinib (Rux; 400 nM) was added to the cultures during the 24 hours of infection. To inhibit phagocytosis, macrophages were pretreated with Cytochalasin D (CytD; 5 μM) for 30 min before addition to organoids. From supernatants of these cultures, cytokine profiles were measured at 24 hours post-infection and plotted as fold-change relative to mock-infected organoids (B). Virus titer data is represented as geometric mean. Statistical differences in the area under the curve (AUC) of viral titers between conditions were assessed using the log-normalized repeated measures one-way ANOVA followed by Tukey posthoc test (* = *p* < 0.05, ** = *p* < 0.01). Scalebar = 50 μM. mФ = macrophages.

### Neutrophils reduce extracellular influenza virus through epithelial-independent mechanisms

The addition of neutrophils to organoid cultures resulted in robust antiviral activity in infected organoids, consistently reducing viral titers across the experiments (**Fig. 4A**). However, despite this reduction, neutrophils did not prevent virus dissemination throughout the organoids (**Fig. 4D,E**), as evidenced by the comparable distribution and intensity of nucleoprotein staining relative to that in infected organoids without neutrophils (**Fig. 4B,C**). Of note, viral replication did not occur in neutrophils (data not shown).

**Figure 4.**
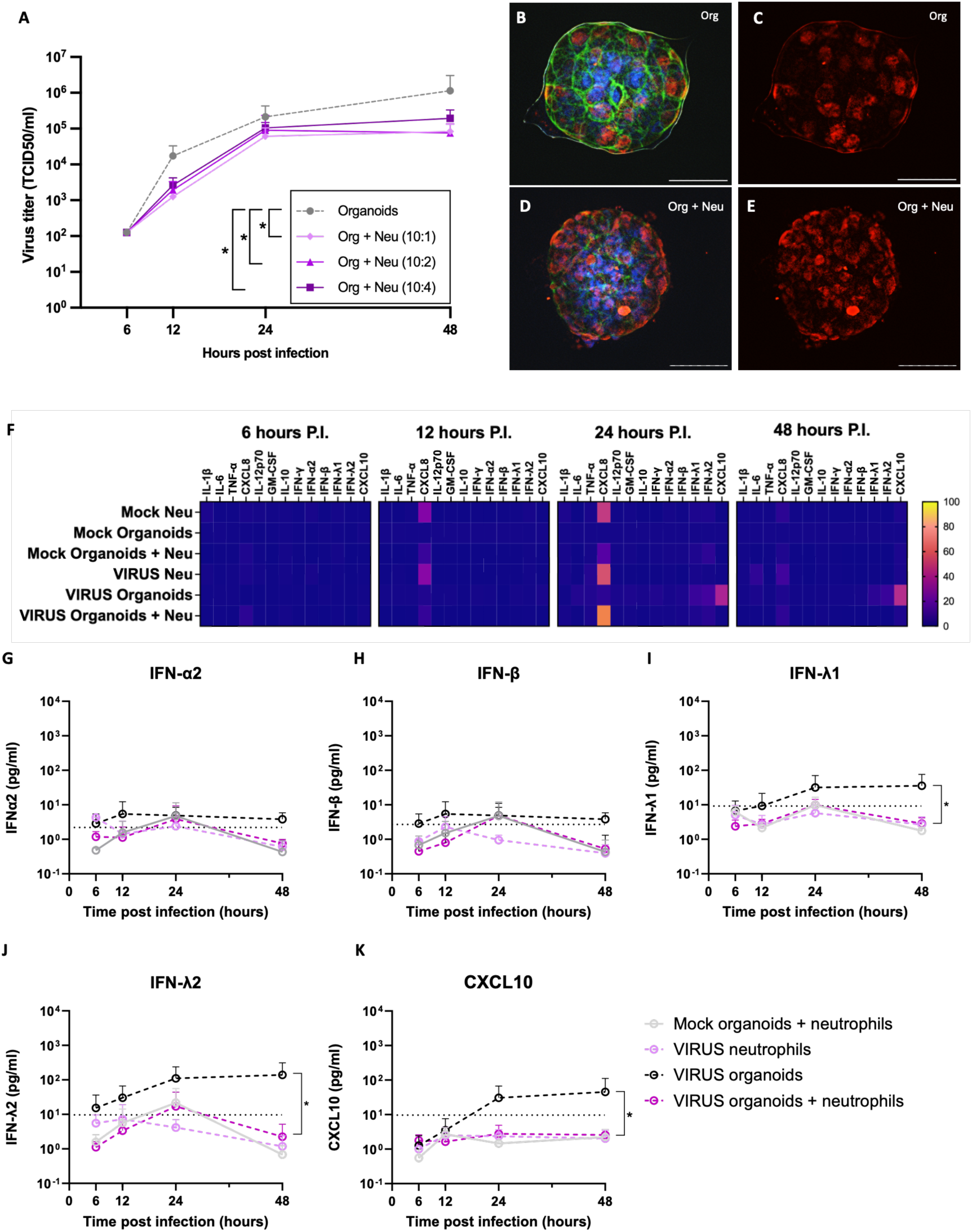
Impact of neutrophils on influenza virus infection in AO airway organoids. Influenza virus titers were quantified from co-cultures of infected organoids (MOI 0.01) and neutrophils (A). At 24 hours post-infection, organoids were stained to visualize influenza virus dissemination. Infected organoids cultured without neutrophils (B, C) or with neutrophils (D, E) were stained for influenza virus nucleoprotein (red), nuclei (DAPI, blue), and filamentous actin (Phalloidin, green). Fluorescent microscopy pictures were taken at 400x magnification. Cytokine levels were measured in culture supernatants at 6, 12, 24, and 48 hours post-infection and plotted as fold-change relative to mock-infected organoids. Specifically, levels of antiviral cytokines IFN-α2 (G), IFN-β (H), IFN-λ1 (I), IFN-λ2 (J), and the IFN-inducible chemokine CXCL10 (K) are shown. Virus titers and cytokine data are represented as geometric mean ± SD and mean ± SD, respectively. Statistical differences in the area under the curve (AUC) of viral titers and cytokine responses between conditions were assessed using the log-normalized repeated measures one-way ANOVA followed by Tukey posthoc test and Friedman test followed by Dunn’s post hoc test, respectively (* = *p* < 0.05). Scalebar = 50 μM. mФ = macrophages; P.I. = post infection.

In both monoculture and co-culture conditions, neutrophils mounted a relatively muted cytokine response, regardless of infection status (**Fig. 4F**). CXCL8 was the only cytokine markedly elevated in the presence of neutrophils (**Suppl. Fig 2B**). Strikingly, the infection-induced secretion of IFN-λ1 (**Fig. 4I**), IFN-λ2 (**Fig. 4J**), and CXCL10 (**Fig. 4K**), typically observed in infected organoids, was reduced when organoids were co-cultured with neutrophils, independent of the number of neutrophils in culture (**Suppl. Fig. 2B**). Taken together, these findings suggest that neutrophils limit extracellular virus through mechanisms independent of epithelial cells, prompting us to investigate whether phagocytosis or trapping of extracellular virus in neutrophil extracellular traps (NETs), indirectly determined by measuring by neutrophil granulation, contributed to this effect.

To assess whether the neutrophil-mediated reduction in influenza virus titers was dependent on viral phagocytosis, neutrophil-organoid cocultures were treated with the phagocytosis inhibitor cytochalasin D. Inhibition of phagocytosis partly limited neutrophil-induced reduction in virus levels in the supernatant, suggesting that neutrophils partly limit infection through phagocytosis (**Fig. 5A**). To determine the level of neutrophil granulation, myeloperoxidase (MPO) levels in culture supernatants were quantified (**Fig. 5B**). Neutrophils markedly increased MPO levels over time, both in the absence and presence of influenza virus. Upon infection, however, MPO concentrations were higher compared to uninfected cocultures at 12 hours and 24 hours post-infection. Importantly, this increase was specific to neutrophil–organoid cocultures and was not observed in organoid cultures alone or in cocultures with macrophages or monocytes (**Suppl. Fig. 3**). Together, these findings suggest that the neutrophil mediated reduction in influenza virus titers is associated with enhanced neutrophil phagocytosis and degranulation.

**Figure 5.**
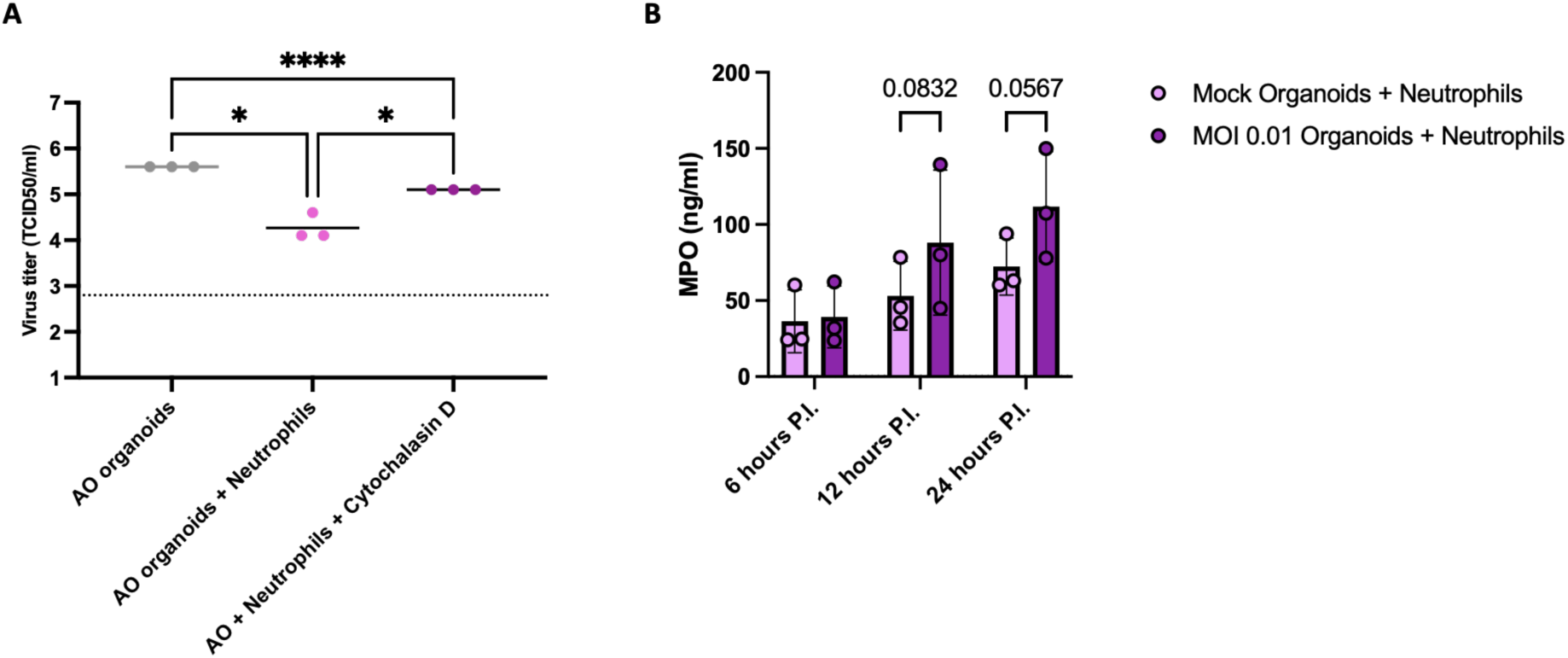
Factors underlying the antiviral effects of neutrophils. Influenza virus titers were quantified from infected AO airway organoid cultures, and co-cultures between neutrophils and infected organoids (A). To inhibit phagocytosis in neutrophils, neutrophils were pretreated with Cytochalasin D (5 μM) for 30 min, before addition to organoids. To determine neutrophil degranulation, myeloperoxidase levels were determined from the supernatant of mock-infected and infected co-cultures between AO airway organoids and neutrophils at 6, 12 and 24 hours post-infection (B). Data of virus titers and MPO levels are represented as geometric mean ± SD and mean ± SD, respectively. Statistical differences in virus titers were determined with log-normalized one-way ANOVA followed by Tukey post-hoc test, whereas repeated-measures two-way ANOVA followed by Bonferroni post-hoc test was used for analyzing differences in MPO levels between mock and MOI conditions (* = *p* < 0.05, ** = *p* < 0.01).*** = *p* < 0.001).

### Monocytes moderately limit viral replication in influenza virus-infected organoids

Co-culturing organoids with monocytes did not significantly alter viral replication kinetics, as virus titers remained comparable to those observed in organoids infected in the absence of monocytes over a 48-hour period (**Fig. 6A**). Moreover, monocytes did not prevent viral replication in organoids (**Fig. 6D,E**).

**Figure 6.**
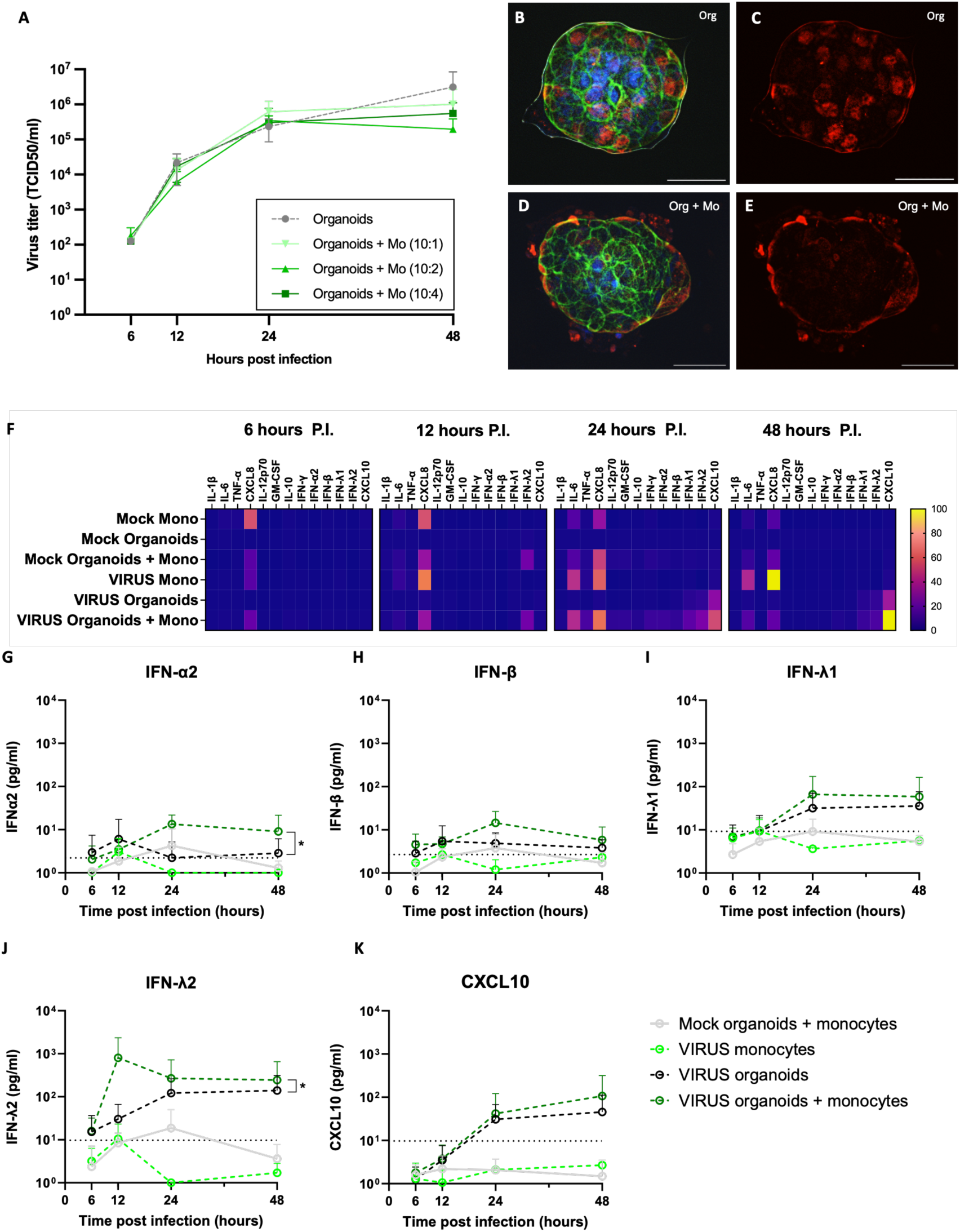
Impact of monocytes on influenza virus infection in AO airway organoids. Influenza virus titers were quantified from co-cultures of infected organoids (MOI 0.01) with monocytes (A). At 24 hours post-infection, organoids were stained to visualize influenza virus dissemination. Infected organoids cultured without monocytes (B, C) or with monocytes (D, E) were stained for influenza virus nucleoprotein (red), nuclei (DAPI, blue), and filamentous actin (Phalloidin, green). Cytokine levels were measured in culture supernatants at 6, 12, 24, and 48 hours post-infection and plotted as fold-change relative to mock-infected organoids. Specifically, levels of antiviral cytokines IFN-α2 (G), IFN-β (H), IFN-λ1 (I), IFN-λ2 (J), and the IFN-inducible chemokine CXCL10 (K) are shown. Fluorescent microscopy pictures were taken at 400x magnification. Virus titers and cytokine data are represented as geometric mean ± SD and mean± SD, respectively. Statistical differences in the area under the curve (AUC) of viral titers and cytokine responses between MOI conditions were assessed using the Friedman test followed by Dunn’s post hoc test (* = *p* < 0.05). Scalebar = 50 μM. mФ = macrophages; P.I. = post infection. Scalebar = 50 μM. mФ = macrophages; P.I. = post infection.

Monocytes, whether in monoculture or co-culture, triggered a rapid and sustained increase in CXCL8 secretion, which persisted up to 48 hours post-infection (**Fig. 6F**). Co-culturing monocytes with infected organoids led to an increase of the cytokines IL-6, TNF-α and IFN-γ which are related to pro-inflammatory macrophages. However, the levels of IL-10, linked to anti-inflammatory macrophages, did not increase upon infection (**Fig. 4F**). Additionally, moderate levels of IFN-α (**Fig. 6G**) were detected in infected monocyte-organoid co-cultures, while secretion of IFN-λ2 (**Fig. 6J**) was more prominently increased than observed for infected organoids in the absence of monocytes. Furthermore, infected monocyte-organoid co-cultures also showed higher levels of IFN-β (**Fig. 6H**) and IFN-λ1 (**Fig. 6I**), yet not at higher levels than infected organoids monocultures. Notably, this cytokine secretion profile was absent in infected monocyte or organoid monocultures and was less pronounced when monocyte numbers were reduced (**Suppl. Fig. 2C**). Collectively, these findings indicate that monocytes contribute to the induction of antiviral cytokines, albeit at moderate levels that may explain their relatively limited capacity to suppress viral replication in epithelial cells, as opposed to the more pronounced effects observed with macrophages.

## Discussion

In this study, we exploited an immune-competent apical-out (AO) human airway organoid model to dissect how distinct innate immune cell types regulate early influenza A virus infection at the respiratory epithelium. Our data demonstrate that macrophages restrict epithelial influenza virus replication primarily through type I and type III interferon (IFN) signaling, whereas neutrophils reduce viral clearance through extraepithelial mechanisms. Monocytes did not reduce virus titers. With their potential to limit viral replication in the epithelium, our study suggests macrophages as cellular targets for future strategies to prevent influenza virus infection.

To dissect the antiviral roles of immune cells during epithelial influenza virus infection, we leverage AO airway organoids. Our findings demonstrate that such organoids recapitulate key features of the human upper airway epithelium, including cellular composition, sialic acid receptors distribution, and susceptibility to productive influenza A virus infection (19, 26). Upon infection, the organoids mounted an epithelial-intrinsic cytokine response characterized mainly by the induction of CXCL8, CXCL10, and type III interferons. In contrast to the human upper airway mucosa, where influenza infection rapidly induces a broader pro-inflammatory cytokine milieu including IL-1β, TNF-α, IL-6, CXCL8, CXCL10 and type I interferons (27–29), this restricted response underscores the importance of epithelial–immune cell interactions in shaping the full mucosal cytokine landscape during influenza virus infection. Extending these observations, we systematically compared infection dynamics across different multiplicities of infection (MOIs). Surprisingly, infection at a low MOI (0.001) yielded the highest viral titers over time, whereas higher MOIs triggered stronger cytokine responses but limited replication. These observations suggest that low-dose infections (MOI 0.001) may support more sustained viral replication over time. A lower MOI likely results in infection of a limited number of cells, allowing the virus to propagate through multiple rounds of replication. In contrast, high MOIs may cause widespread infection early on, leading to rapid cytopathic effects and structural disintegration of the organoids, thereby limiting further replication (30). Another possible explanation is the increased generation of defective interfering particles at high MOI, which can suppress productive viral replication over time (31).

Among the innate immune cells tested, macrophages exerted the most pronounced antiviral effect that correlated with robust secretion of type I and III IFNs. By pharmacologically uncoupling IFN signaling and phagocytosis, we demonstrate that inhibition of JAK–STAT signaling consistently abrogates macrophage-mediated viral control, establishing IFN induction as an essential antiviral mechanism in this system. In contrast, blockade of phagocytosis with cytochalasin D unexpectedly enhanced IFN production and resulted in further reductions in viral titers. Although counterintuitive, this observation suggests that active phagocytosis can suppress macrophage IFN responses during epithelial influenza infection. Phagocytosis, particularly of apoptotic or infected epithelial cells, is increasingly recognized as an immunoregulatory process of macrophages that promotes resolution of inflammation and limits excessive cytokine production (10, 32). Engagement of such pathways can activate negative feedback mechanisms that dampen interferon signaling. In our system, blocking phagocytosis appeared to release this restraint, allowing sustained pathogen sensing through TLR3, TLR7, and RIG-I, resulting in amplified type I and III IFN production (33,34). The dampening of the cytochalasin D–induced antiviral effect upon concomitant JAK1/2 inhibition suggests that the observed viral control under phagocytosis blockade is mediated indirectly via IFN signaling rather than by reduced uptake of viral particles. Together, our findings reveal that macrophage phagocytosis and interferon production are not necessarily synergistic antiviral pathways but can function in opposition during influenza infection of the airway epithelium.

The observed cytokine responses align with *in vivo* and other *in vitro* studies showing that human macrophages produce type I IFN (IFN-α, IFN-β), type III IFN (IFN-λ1, IFN-λ2), TNFα and IL-6 (35–37). Moreover, the increase in IFN-γ, TNF-α, IL-6, and CXCL10, cytokines typically associated with pro-inflammatory macrophages, suggests that viral infection in organoids promotes differentiation of unpolarized macrophages toward a pro-inflammatory antiviral phenotype rather than an IL-10-producing anti-inflammatory phenotype (32,35). Nevertheless, some degree of anti-inflammatory polarization appears to occur during influenza infection as shown by the persistent levels of IL-10 in the presence of macrophages. These findings align with animal studies showing that pro-inflammatory macrophages dominate the acute phase of infection, whereas anti-inflammatory macrophages contribute to resolution and protection against excessive tissue damage in the lungs (32,37,38). Moreover, elevated cytokine levels were detected only in co-cultures where infected organoids were in direct contact with macrophages and not in other conditions. This finding indicates that direct macrophage–epithelial interactions are critical for robust cytokine induction. This observation is supported by previous work demonstrating that physical contact between macrophages and epithelial cells is required for effective viral control (39), whereas cytokine secretion by virus-activated macrophages in the absence of cell–cell contact is insufficient to suppress infection. Together, these findings identify macrophages as a potent therapeutic target for restricting viral infection at the epithelium.

In contrast to macrophages, neutrophils, and monocytes appear less suitable as primary therapeutic targets. Neutrophils markedly reduced virus titers, they were insufficient to prevent viral replication in organoids. In fact, the addition of neutrophils markedly reduced IFN induction, most likely through cytokine or virus scavenging (40), thereby leaving the epithelium more exposed to infection. Instead, neutrophil-mediated viral control appeared to rely primarily on phagocytosis and degranulation, consistent with mechanisms such as neutrophil extracellular trap formation (41). In contrast, monocytes did not prevent influenza virus replication in organoids. Their low ability to regulate epithelial infection may be explained by their less prominent cytokine profile compared to that of macrophages, which is characterized by the temporary activation of IL-6, TNF-α, and type I/III interferons. Monocytes probably need to first differentiate into macrophages or dendritic cells, before they contribute strongly to antiviral effects (10). Future studies should therefore investigate monocyte differentiation dynamics in time, as well as the delayed contributions of monocytes to antiviral defense and immunoregulation.

Several limitations of the current study point to important future directions. First, the observation window was restricted to 48 hours and therefore primarily captured early infection events. Extending cultures over longer periods, particularly in macrophage and monocyte cultures, would enable investigation of later phases of infection, including epithelial repair, immune cell adaptation, and resolution of inflammation. Second, the use of a single epithelial donor for organoid development in co-culture precludes conclusions regarding inter-individual variability. Given our hypothesis that immune cells play a dominant role in controlling viral replication, we deliberately included immune cells from multiple donors to capture immune-intrinsic variability. Future studies should also assess epithelial donor variability and its impact on immune-epithelial interactions. Moreover, pairing epithelial and immune cells from the same donor would allow interrogation of autologous immune-epithelial networks and help disentangle epithelial-intrinsic versus immune-intrinsic contributions to antiviral defense.

In summary, this study reveals that macrophages control early influenza virus infection of the human airway epithelium predominantly through the induction of type I and type III interferons, rather than through direct phagocytic clearance of virus particles. By functionally separating these macrophage effector pathways in a human-relevant epithelial context, we uncover an unexpected immunoregulatory role for phagocytosis in dampening antiviral interferon responses. In contrast, neutrophils and monocytes exert distinct and more limited antiviral functions. Together, our findings highlight the complexity of epithelial-immune cell interactions during influenza infection and demonstrate how controlled immune-organoid cocultures can be exploited to dissect mechanisms of mucosal antiviral defense. This platform may prove valuable for identifying correlates of protection and informing strategies aimed at enhancing antiviral immunity at the respiratory mucosa.

## Supplementary Figures

**Supplementary Figure 1.**
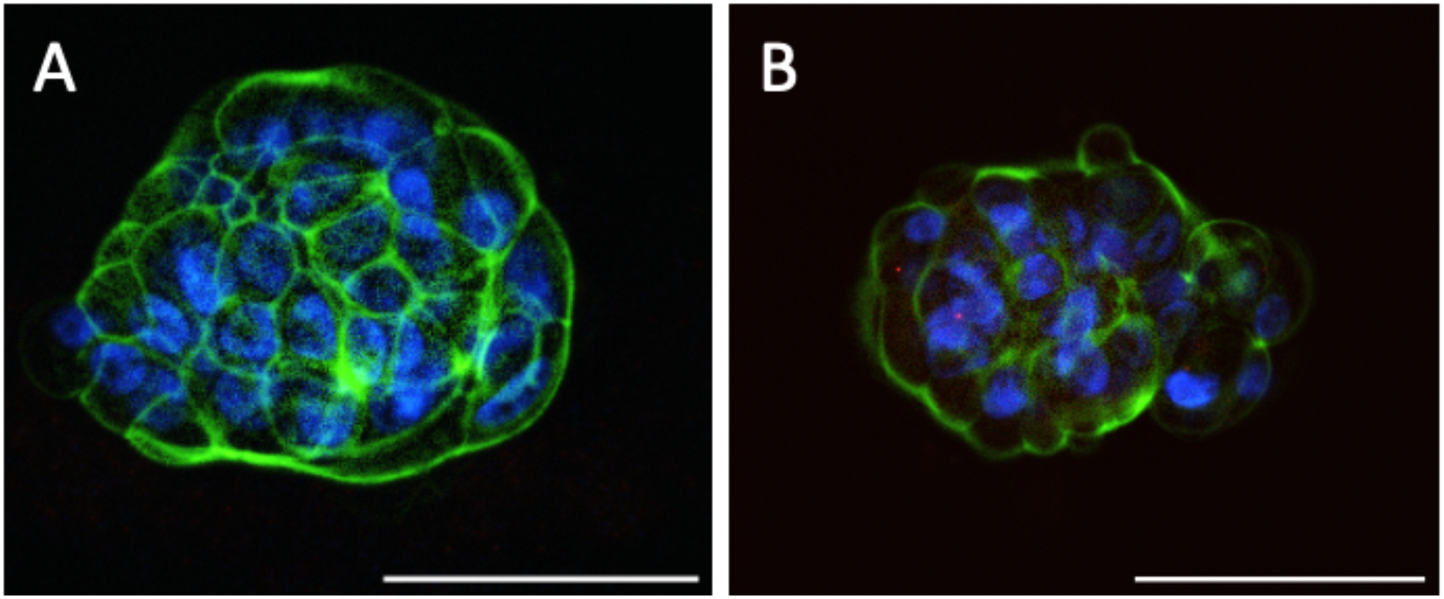
Immunofluorescent phenotyping of lung epithelial cell types in apical out airway organoids. The organoids were fluorescently stained to detect the presence of club cells (A; CC10) or alveolar type 2 epithelial cells (B; Pro-SPC = Pro surfactant protein. Blue dye indicates nuclear staining (DAPI). Green dye indicates filamentous actin (Phalloidin). Fluorescent microscopy pictures were taken at 400x magnification. Scale bar = 50 µM.

**Supplementary Figure 2.**
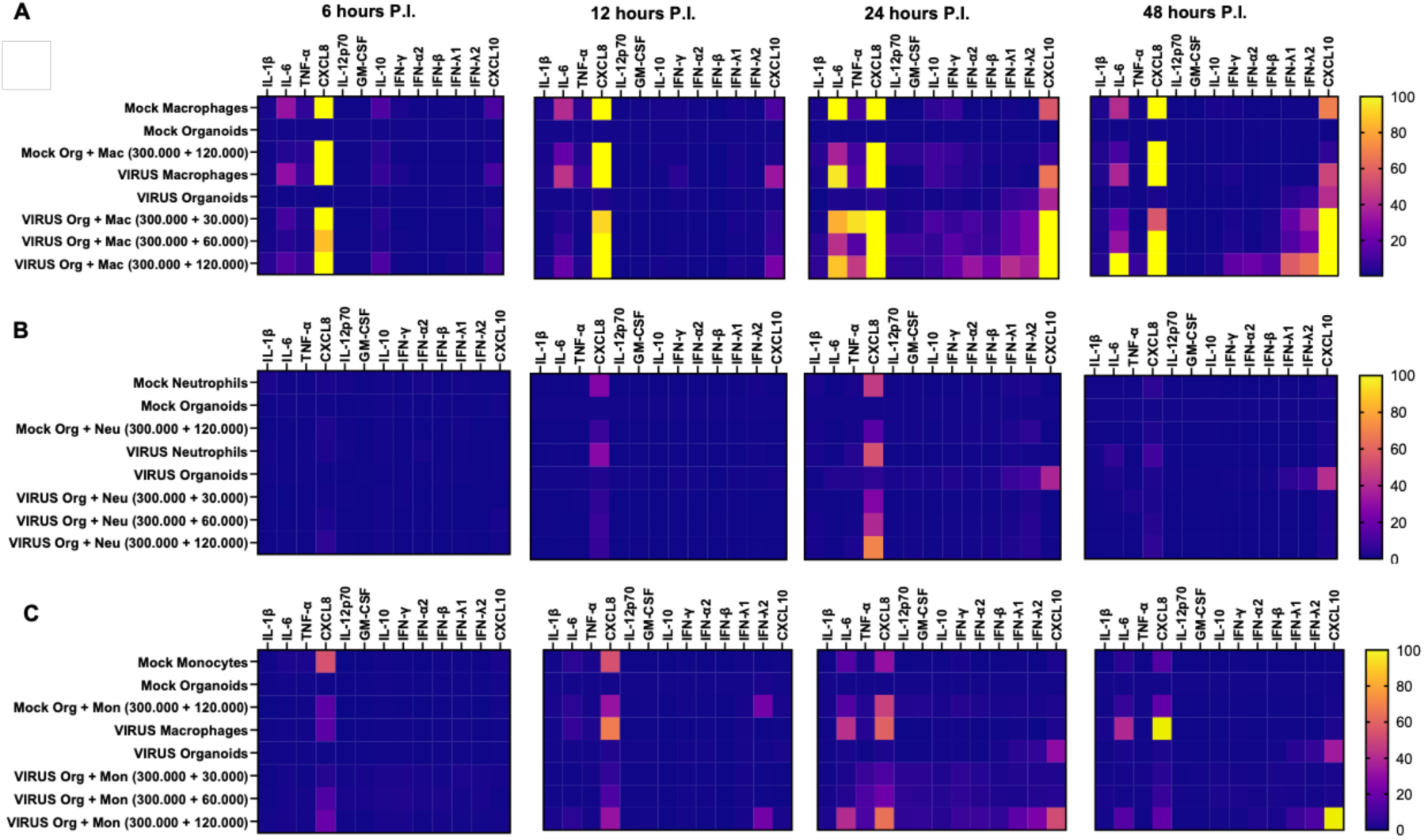
Virus-induced cytokine levels in co-cultures of tracheal organoids and neutrophils (A), macrophages (B), or monocytes (C). Co-culture conditions were standardized by using a consistent number of epithelial cells (300,000), while varying the number of immune cells added (30,000, 60,000, and 120,000). This resulted in epithelial cell-to-monocyte ratios of 10:1, 10:2, and 10:4, respectively. The heatmaps illustrate the fold-change in cytokine levels at each time point relative to cytokine levels at the 6-hour time point. PI = post infection.

**Supplementary Figure 3.**
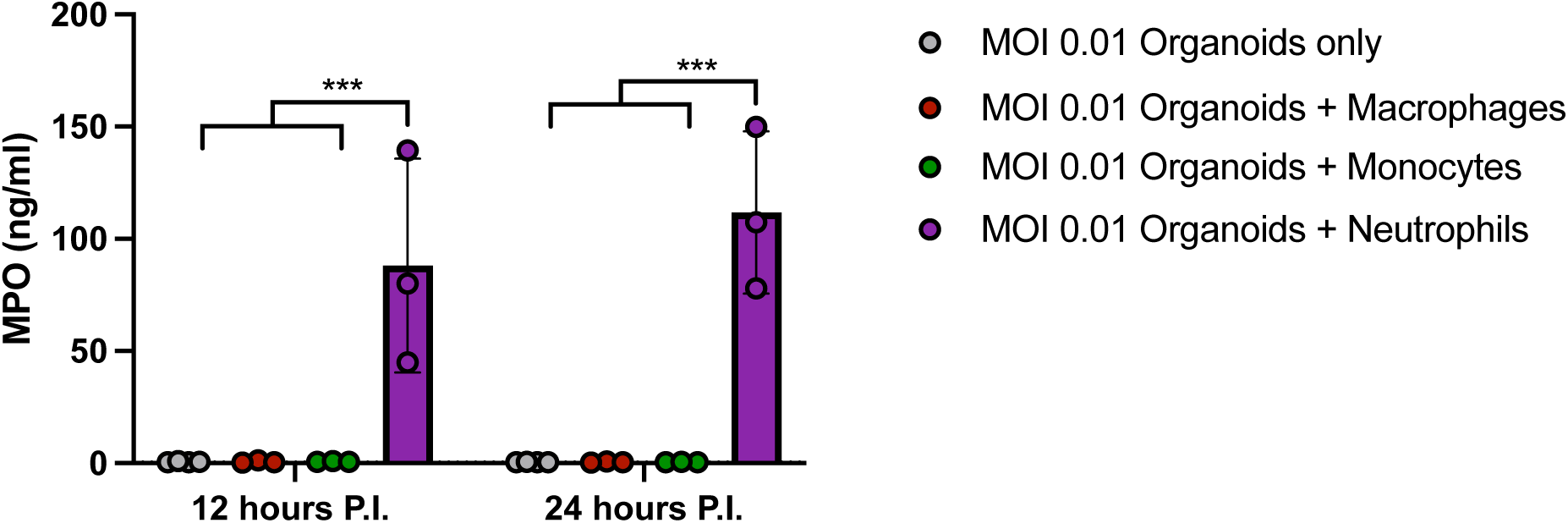
MPO levels of virus-infected AO airway organoids and co-cultures. MPO levels in supernatants from co-cultures were measured at 24 hours post infection.

## References

[1] Influenza (seasonal), World Health Organisation, 2025 feb, https://www.who.int/news-room/fact-sheets/detail/influenza-(seasonal)

[2] GBD 2019 Chronic Respiratory Diseases Collaborators. Global burden of chronic respiratory diseases and risk factors, 1990-2019: an update from the Global Burden of Disease Study 2019. EClinicalMedicine. 2023 May;59:101936.

[3] Bean R, Giurgea LT, Han A, Czajkowski L, Cervantes-Medina A, Gouzoulis M, Mateja A, Hunsberger S, Reed S, Athota R, Baus HA, Kash JC, Park J, Taubenberger JK, Memoli MJ. Mucosal correlates of protection after influenza viral challenge of vaccinated and unvaccinated healthy volunteers. mBio. 2024 Feb 14;15(2):e0237223.

[4] Morens DM, Taubenberger JK, Fauci AS. Rethinking next-generation vaccines for coronaviruses, influenzaviruses, and other respiratory viruses. Cell Host Microbe. 2023 Jan 11;31(1):146–157.

[5] Matrosovich MN, Matrosovich TY, Gray T, Roberts NA, Klenk HD. Human and avian influenza viruses target different cell types in cultures of human airway epithelium. Proc Natl Acad Sci U S A. 2004 Mar 30;101(13):4620–4.

[6] Latino I, Gonzales SF, Spatio-temporal profile of innate inflammatory cells and mediators during influenza virus infection, Current Opion in Physiology. 2021 Feb;19:175–186.

[7] Márquez-Bandala AH, Gutierrez-Xicotencatl L, Esquivel-Guadarrama F. Pathogenesis Induced by Influenza Virus Infection: Role of the Early Events of the Infection and the Innate Immune Response. Viruses. 2025 May 12;17(5):694.

[8] Gao KM, Derr AG, Guo Z, Nündel K, Marshak-Rothstein A, Finberg RW, Wang JP. Human nasal wash RNA-Seq reveals distinct cell-specific innate immune responses in influenza versus SARS-CoV-2. JCI Insight. 2021 Nov 22;6(22):e152288.

[9] Ballesteros I, Rubio-Ponce A, Genua M, Lusito E, Kwok I, Fernández-Calvo G, Khoyratty TE, van Grinsven E, González-Hernández S, Nicolás-Ávila JÁ, Vicanolo T, Maccataio A, Benguría A, Li JL, Adrover JM, Aroca-Crevillen A, Quintana JA, Martín-Salamanca S, Mayo F, Ascher S, Barbiera G, Soehnlein O, Gunzer M, Ginhoux F, Sánchez-Cabo F, Nistal-Villán E, Schulz C, Dopazo A, Reinhardt C, Udalova IA, Ng LG, Ostuni R, Hidalgo A. Co-option of Neutrophil Fates by Tissue Environments. Cell. 2020 Nov 25;183(5):1282–1297.e18.

[10] Ruscitti C, Radermecker C, Marichal T. Journey of monocytes and macrophages upon influenza A virus infection. Curr Opin Virol. 2024 Jun;66:101409.

[11] Gong HH, Worley MJ, Carver KA, Godin CJ, Deng JC. Deficient neutrophil responses early in influenza infection promote viral replication and pulmonary inflammation. PLoS Pathog. 2025 Jan 17;21(1):e1012449.

[12] Chu JTS, Lamers MM. Organoids in virology. Npj Viruses. 2024 Feb 7;2(1):5.

[13] van der Vaart J, Lamers MM, Haagmans BL, Clevers H. Advancing lung organoids for COVID-19 research. Dis Model Mech. 2021 Jun 1;14(6):dmm049060.

[14] Joo H, Min S, Cho SW. Advanced lung organoids for respiratory system and pulmonary disease modeling. J Tissue Eng. 2024 Feb 22;15:20417314241232502.

[15] Ekanger CT, Zhou F, Bohan D, Lotsberg ML, Ramnefjell M, Hoareau L, Røsland GV, Lu N, Aanerud M, Gärtner F, Salminen PR, Bentsen M, Halvorsen T, Ræder H, Akslen LA, Langeland N, Cox R, Maury W, Stuhr LEB, Lorens JB, Engelsen AST. Human Organotypic Airway and Lung Organoid Cells of Bronchiolar and Alveolar Differentiation Are Permissive to Infection by Influenza and SARS-CoV-2 Respiratory Virus. Front Cell Infect Microbiol. 2022 Mar 14;12:841447.

[16] Svensson L, Nordgren J, Lundkvist Å, Hagbom M. Recent Advances in Nose and Lung Organoid Models for Respiratory Viral Research. Viruses. 2025 Feb 28;17(3):349.

[17] Sachs N, Papaspyropoulos A, Zomer-van Ommen DD, Heo I, Böttinger L, Klay D, Weeber F, Huelsz-Prince G, Iakobachvili N, Amatngalim GD, de Ligt J, van Hoeck A, Proost N, Viveen MC, Lyubimova A, Teeven L, Derakhshan S, Korving J, Begthel H, Dekkers JF, Kumawat K, Ramos E, van Oosterhout MF, Offerhaus GJ, Wiener DJ, Olimpio EP, Dijkstra KK, Smit EF, van der Linden M, Jaksani S, van de Ven M, Jonkers J, Rios AC, Voest EE, van Moorsel CH, van der Ent CK, Cuppen E, van Oudenaarden A, Coenjaerts FE, Meyaard L, Bont LJ, Peters PJ, Tans SJ, van Zon JS, Boj SF, Vries RG, Beekman JM, Clevers H. Long-term expanding human airway organoids for disease modeling. EMBO J. 2019 Feb 15;38(4):e100300.

[18] Watanabe M, Buth JE, Vishlaghi N, de la Torre-Ubieta L, Taxidis J, Khakh BS, Coppola G, Pearson CA, Yamauchi K, Gong D, Dai X, Damoiseaux R, Aliyari R, Liebscher S, Schenke-Layland K, Caneda C, Huang EJ, Zhang Y, Cheng G, Geschwind DH, Golshani P, Sun R, Novitch BG. Self-Organized Cerebral Organoids with Human-Specific Features Predict Effective Drugs to Combat Zika Virus Infection. Cell Rep. 2017 Oct 10;21(2):517–532.

[19] Stroulios G, Brown T, Moreni G, Kondro D, Dei A, Eaves A, Louis S, Hou J, Chang W, Pajkrt D, Wolthers KC, Sridhar A, Simmini S. Apical-out airway organoids as a platform for studying viral infections and screening for antiviral drugs. Sci Rep. 2022 May 10;12(1):7673.

[20] preprint shoran gong PMN paper

[21] Tapia-Calle G, Stoel M, de Vries-Idema J, Huckriede A. Distinctive Responses in an In Vitro Human Dendritic Cell-Based System upon Stimulation with Different Influenza Vaccine Formulations. Vaccines (Basel). 2017 Aug 9;5(3):21.

[22] Horvathova L, Kinderman P, Janzen T, Hoogeboom BN, Weissing FJ, van Montfoort N, Daemen T, Bhatt DK. Alphavirus replicons encoding IFN-γ enhance cancer virotherapy by overcoming macrophage-mediated suppression. iScience. 2025 Sep 10;28(10):113545.

[23] Schulz D, Severin Y, Zanotelli VRT, Bodenmiller B. In-Depth Characterization of Monocyte-Derived Macrophages using a Mass Cytometry-Based Phagocytosis Assay. Sci Rep. 2019 Feb 13;9(1):1925.

[24] Beukema M, Gong S, Al-Jaawni K, de Vries-Idema JJ, Krammer F, Zhou F, Cox RJ, Huckriede A. Prolonging the delivery of influenza virus vaccine improves the quantity and quality of the induced immune responses in mice. Front Immunol. 2023 Oct 5;14:1249902.

[25] C. Spearman, the Method of ‘Right and Wrong Cases’ (‘Constant Stimuli’) Without Gauss’S Formulae, Br. J. Psychol. 1908; 1904–1920 (2):227–242

[26] Davis JD, Wypych TP. Cellular and functional heterogeneity of the airway epithelium. Mucosal Immunol. 2021 Sep;14(5):978–990.

[27] Bachert C, van Kempen MJ, Höpken K, Holtappels G, Wagenmann M. Elevated levels of myeloperoxidase, pro-inflammatory cytokines and chemokines in naturally acquired upper respiratory tract infections. Eur Arch Otorhinolaryngol. 2001 Oct;258(8):406–12.

[28] Julkunen I, Melén K, Nyqvist M, Pirhonen J, Sareneva T, Matikainen S. Inflammatory responses in influenza A virus infection. Vaccine. 2000 Dec 8;19 Suppl 1:S32–7.

[29] Hayden FG, Fritz R, Lobo MC, Alvord W, Strober W, Straus SE. Local and systemic cytokine responses during experimental human influenza A virus infection. Relation to symptom formation and host defense. J Clin Invest. 1998 Feb 1;101(3):643–9.

[30] Ekanger CT, Dinesh Kumar N, Koutstaal RW, Zhou F, Beukema M, Waldock J, Jochems SP, Mulder N, van Els CACM, Engelhardt OG, Mantel N, Buno KP, Brokstad KA, Engelsen AST, Cox RJ, Melgert BN, Huckriede ALW, van Kasteren PB. Comparison of air-liquid interface transwell and airway organoid models for human respiratory virus infection studies. Front Immunol. 2025 Feb 6;16:1532144.

[31] Bora, M., Manu, M. & Karki, M. Exploring the mechanisms of interference, persistence and antiviral potential of defective interfering particles. Discov. Viruses. 2025 Jan; 2(1).

[32] Cline TD, Beck D, Bianchini E. Influenza virus replication in macrophages: balancing protection and pathogenesis. J Gen Virol. 2017 Oct;98(10):2401–2412.

[33] Rothlin CV, Ghosh S, Zuniga EI, Oldstone MB, Lemke G. TAM receptors are pleiotropic inhibitors of the innate immune response. Cell. 2007 Dec 14;131(6):1124–36. doi: 10.1016/j.cell.2007.10.034. PMID: 18083102.

[34] A-Gonzalez N, Bensinger SJ, Hong C, Beceiro S, Bradley MN, Zelcer N, Deniz J, Ramirez C, Díaz M, Gallardo G, de Galarreta CR, Salazar J, Lopez F, Edwards P, Parks J, Andujar M, Tontonoz P, Castrillo A. Apoptotic cells promote their own clearance and immune tolerance through activation of the nuclear receptor LXR. Immunity. 2009 Aug 21;31(2):245–58.

[35] Li H, Wang A, Zhang Y, Wei F. Diverse roles of lung macrophages in the immune response to influenza A virus. Front Microbiol. 2023 Sep 13;14:1260543.

[36] Dutry I, Li J, Hung Li P, Bruzzone R, Malik Peiris JS, Jaume M, The effects of macrophage polarity on influenza virus replication and innate immune responses. J Clin. & Cell Immunology. 2015 Feb 23; 6:1

[37] Cole SL, Dunning J, Kok WL, Benam KH, Benlahrech A, Repapi E, Martinez FO, Drumright L, Powell TJ, Bennett M, Elderfield R, Thomas C; MOSAIC investigators; Dong T, McCauley J, Liew FY, Taylor S, Zambon M, Barclay W, Cerundolo V, Openshaw PJ, McMichael AJ, Ho LP. M1-like monocytes are a major immunological determinant of severity in previously healthy adults with life-threatening influenza. JCI Insight. 2017 Apr 6;2(7):e91868.

[38] Wu W, Zhang W, Alexandar JS, Booth JL, Miller CA, Xu C, Metcalf JP. RIG-I agonist SLR10 promotes macrophage M1 polarization during influenza virus infection. Front Immunol. 2023 Jul 5;14:1177624.

[39] Barreto-Duran E, Szczepański A, Gałuszka-Bulaga A, Surmiak M, Siedlar M, Sanak M, Rajfur Z, Milewska A, Lenart M, Pyrć K. The interplay between the airway epithelium and tissue macrophages during the SARS-CoV-2 infection. Front Immunol. 2022 Oct 6;13:991991.

[40] Siwicki M, Kubes P. Neutrophils in host defense, healing, and hypersensitivity: Dynamic cells within a dynamic host. J Allergy Clin Immunol. 2023 Mar;151(3):634–655.

[41] Gong HH, Worley MJ, Carver KA, Godin CJ, Deng JC. Deficient neutrophil responses early in influenza infection promote viral replication and pulmonary inflammation. PLoS Pathog. 2025 Jan 17;21(1):e1012449.

