## Supplementary figures and images for "Immune-competent apical-out airway organoids reveal distinct antiviral strategies of macrophages, neutrophils, and monocytes during influenza infection"

### Supplementary Figure 1

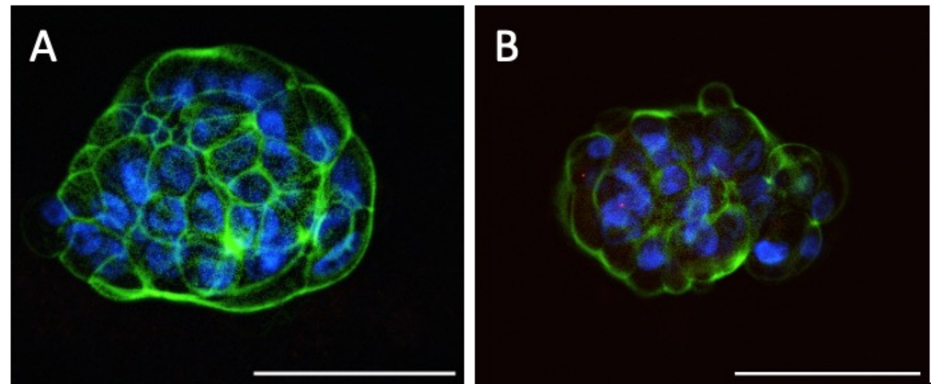

### Supplementary Figure 2

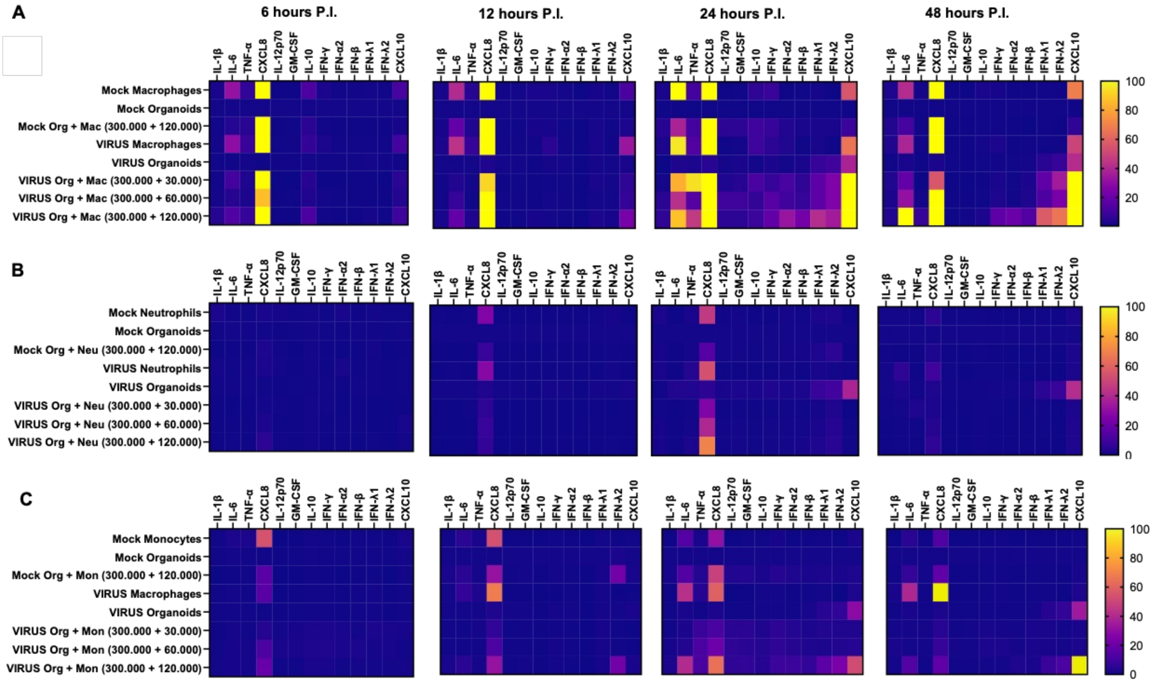

### Supplementary Figure 3

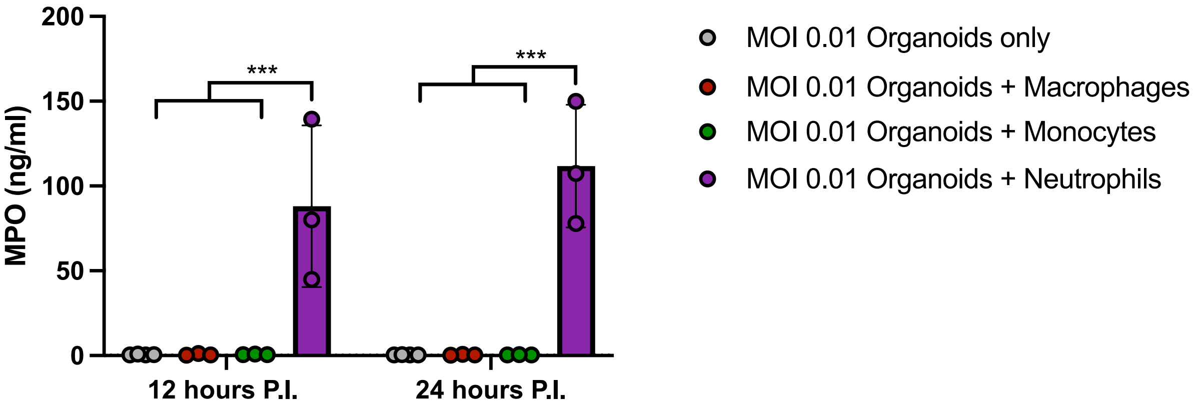
